# Additive manufactured scaffolds for bone tissue engineering: physical characterization of thermoplastic composites with functional fillers

**DOI:** 10.1101/2021.03.23.436548

**Authors:** Ravi Sinha, Alberto Sanchez, Maria Camara-Torres, Iñigo Calderon Uriszar-Aldaca, Andrea Roberto Calore, Jules Harings, Ambra Gambardella, Lucia Ciccarelli, Veronica Vanzanella, Michele Sisani, Marco Scatto, Rune Wendelbo, Sergio Perez, Sara Villanueva, Amaia Matanza, Alessandro Patelli, Nino Grizzuti, Carlos Mota, Lorenzo Moroni

## Abstract

Thermoplastic polymer – filler composites are excellent materials for bone tissue engineering (TE) scaffolds, combining the functionality of fillers with suitable load bearing ability, biodegradability, and additive manufacturing (AM) compatibility of the polymer. Two key determinants of their utility are their rheological behavior in the molten state, determining AM processability, and their mechanical load-bearing properties. We report here the characterization of both these physical properties for four bone TE relevant composite formulations with poly(ethylene oxide terephthalate) / poly(butylene terephthalate (PEOT/PBT) as a base polymer, which is often used to fabricate TE scaffolds. The fillers used were reduced graphene oxide (rGO), hydroxyapatite (HA), gentamycin intercalated in zirconium phosphate (ZrP-GTM) and ciprofloxacin intercalated in MgAl layered double hydroxide (MgAl-CFX). The rheological assessment showed that generally the viscous behavior dominated the elastic behavior (G’’ > G’) for the studied composites, at empirically determined extrusion temperatures. Coupled rheological-thermal characterization of ZrP-GTM and HA composites showed that the fillers increased the solidification temperatures of the polymer melts during cooling. Both these findings have implications for the required extrusion temperatures and bonding between layers. Mechanical tests showed that the fillers generally made the polymer stiffer but more brittle in proportion to the filler fractions. Furthermore, the elastic moduli of scaffolds did not directly correlate with the corresponding bulk material properties, implying composite-specific AM processing effects on the mechanical properties. Lastly, we show computational models to predict multi-material scaffold elastic moduli using measured single material scaffold and bulk moduli. The reported characterizations are essential for assessing the AM processability and ultimately the suitability of the manufactured scaffolds for the envisioned bone regeneration application.

## Introduction

Poly(ethylene oxide terephthalate) / poly(butylene terephthalate (PEOT/PBT) thermoplastic block co-polymers have shown promise as biomaterials for the fabrication of 3D scaffolds for tissue engineering (TE) applications [1-3]. Diverse PEOT/PBT block copolymers have been investigated for a multitude of TE applications spanning from soft tissues, such as skin [4] and neural regeneration [5], to hard skeletal tissue such as bone and cartilage regeneration [6, 7]. These copolymers have a broad range of mechanical properties (modulus in the range 40 – 300 MPa [8]), tunable degradation rate by adjusting the PEOT and PBT ratio [8, 9], are commonly used for the manufacturing of scaffolds using fused deposition modeling (FDM) additive manufacturing (AM) [10, 11], and have demonstrated good performance *in vitro* [12] and *in vivo* [2] for bone regeneration. The formulation 300PEOT55PBT45, consisting of PEOT and PBT in the ratio 55:45 and prepared from a starting molecular weight of PEO of 300 g/mol, has often been investigated for bone TE owing to its intermediate degradation rate and good adhesion to existing bone [9].

The addition of fillers into a thermoplastic matrix has been investigated to further strengthen the overall material properties and the regeneration capacity of the scaffolds post-implantation. Calcium phosphates (CaP) are a good example of these fillers which have shown to improve the bone formation outcome when combined with a PEOT/PBT polymeric matrix [13]. Dispersion of CaP into the polymeric phase has shown mechanical property improvement on scaffolds manufactured with AM [14, 15]. Increasing the fraction of CaPs into a polymer matrix generally makes the composites stiffer than the polymer alone, thereby improving their load bearing abilities, bringing them closer to those of bone (Young’s modulus 0.1-2 GPa for trabecular and 15-20 GPa for cortical bone [16]), which itself has a high mineral content (∼60% w/w) [17]. Thus, increasing the filler loadings in polymers for producing bone TE scaffolds is needed, yet difficult to achieve.

Besides the bone composition inspired choice of CaPs as fillers in TE scaffolds, graphene based materials have also attracted attention due to their favorable properties, such as high surface area, high load bearing, availability of reactive groups for chemical functionalization, and electrical conductivity, that can be beneficial for bone TE [18, 19]. Inclusion of graphene-based materials into polymeric scaffold materials is still in a nascent stage [20], but such composites are under active research.

Another filler category, whose inclusion into bone TE scaffolds is being widely explored, is antibiotics that can be locally released while preventing infections at implant sites, avoiding the need for systemic delivery, thereby bypassing side effects [21]. One mode of antibiotic loading that has been utilized is intercalation into inorganic lamellar fillers, where the active compound is held by electrostatic forces while being shielded from thermal degradation during scaffold manufacture. Examples include gentamycin intercalated in zirconium phosphate (ZrP-GTM) [22, 23] and ciprofloxacin intercalated in magnesium aluminum layered double hydroxide (MgAl-CFX) [22].

The ultimate test of polymers and composites, including the previously highlighted fillers, are their ability to improve bone regeneration and additionally to prevent infection in the case of antibiotic-integrated formulations. However, the characterization of their physical properties should precede the scaffolds manufacturing step and the biological validation. The first key property is the rheological behavior of these biomaterials when the polymer phase is in a molten state, which determines their processability with the chosen fused deposition modeling (FDM) technique. A second key property is the mechanical behavior of the bulk and the manufactured scaffolds, which determine if they can withstand the forces arising in the intended application without undergoing failure. The rheological characterization additionally provides indicators of expected scaffold quality parameters, such as layer bonding or sagging of fibers between filaments, besides helping to choose the extrusion temperature and pressure, i.e. a temperature range where the material is liquid enough to be extruded, but viscous enough to prevent flow without the application of a threshold pressure. Good layer bonding is desirable and can be expected when the melt viscous behavior dominates the elastic behavior (although not exclusively) [24], and the material does not solidify faster than needed for bonding. However, the same conditions needed for good bonding can lead to the extruded fibers to collapse under their weight (sagging), being more fluid and for longer times [25]. Thus, a balance needs to be struck between these two situations by choosing appropriately the processing conditions.

This work reports the extensive rheological and mechanical characterization of PEOT/PBT polymer loaded with HA, rGO, ZrP-GTM or MgAl-CFX fillers – a material library developed for producing bone tissue engineering scaffolds. Furthermore, we report how the measured scaffold and bulk mechanical properties can be used to predict the mechanical properties of multi-material scaffolds using computational modeling. The reported characterization will assist researchers in planning scaffold manufacturing, using desired AM techniques, without extensive empirical testing, and in comparing these materials with other available material choices to meet the required load-bearing needs.

## Materials and Methods

### Thermoplastic polymer base

The polymer used was the PEOT/PBT formulation with PEOT and PBT in the ratio 55:45 (w/w) and PEO molecular weight 300 g/mol. The polymer was provided by the manufacturer Polyvation B.V. (Groningen, The Netherlands) as pellets (Figure 1A) and the final molecular weight was reported in terms of intrinsic viscosity (I.V.) at which the polymerization reaction was stopped. Unless specified, I.V. 0.51 dl/g was used. An additional material with I.V. of 0.76 dl/g was also tested for its potential to reduce the brittleness of highly loaded HA composites.

**Figure 1.**
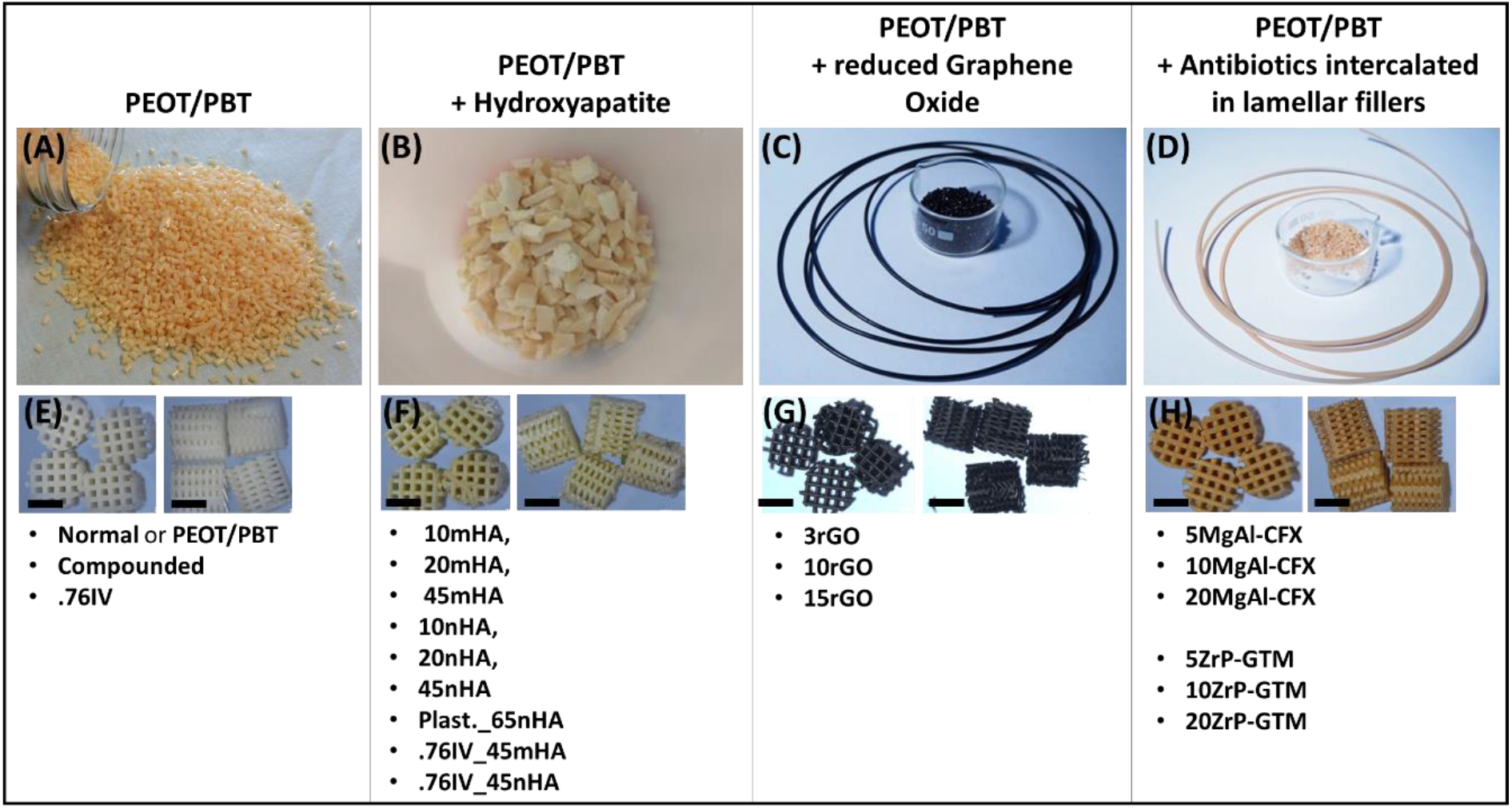
Appearance of the polymer and the composites used (A-D), appearance of representative scaffolds used for mechanical testing (E-H) and list of the variations of each material tested (acronyms used are described in supplementary table S1). Scale bars 2.5 mm.

### HA loaded composite preparation using solvent blending

Hydroxyapatite micro-particles (Nanoxim.HAp200, Fluidinova, 5 ± 1 μm), hereafter referred to as mHA, or nano-particles (Sigma Aldrich, ≤ 200 nm), hereafter referred to as nHA, were loaded into the PEOT/PBT using solvent blending. The polymer was dissolved in chloroform (Scharlab, chloroform to composite ratio 85:15 by weight) under stirring for 45 minutes and the HA particles were mixed in this solution under constant stirring for an additional 15 minutes. The mixture was subsequently precipitated with an excess of a polymer non-solvent (diethyl ether, Scharlab) under stirring. The composite was dried at room temperature, followed by a drying step at 60 °C or 90 °C, both done after decanting the non-solvent. The drying temperature was increased for later batches to further improve solvent removal, but did not show distinguishable effects on the viscosities, hence measurements using the older batch were not repeated. The composites were prepared primarily using 0.51 dl/g I.V. PEOT/PBT. For the high (45%) HA filler loading, samples were also prepared using the 0.76 dl/g I.V. PEOT/PBT in order to reduce brittleness of the composites. The solvent blended composites were compacted into sheets and cut into small pieces for further AM processing and the same form was used for the other reported characterizations (Figure 1B).

### Enhanced HA filler loading using plasticizer

Citric acid ester derivatives were used as plasticizers, due to their expected non-toxicity based on established use in food contact products and reported lower toxicity compared to conventional phthalate plasticizers [26]. The tested plasticizers were: triethyl citrate, acetyltriethyl citrate, tri-n-butyl citrate, acetyltri-n-butyl citrate, acetyltri-n-hexyl citrate, n-butyryltri-n-hexyl citrate (all from the supplier Vertellus). Solvent blending lead to poor mixing between the polymer and the plasticizer, so melt compounding was used. A Brabender Plastograph EC plus kneading mixer was used and mixing was performed at 150 °C, 30 rpm rotation speed, for 15 minutes. The polymer was added first and plasticizer and filler were added after 3.5 and 5 minutes of mixing respectively.

### Graphene and antibiotics composite preparation using melt compounding

The rGO, MgAl-CFX and ZrP-GTM fillers were dispersed into the PEOT/PBT polymer using melt compounding. rGO was obtained from the manufacturer Abalonyx and consisted of rGO prepared by a modified Hummer’s method [27], followed by thermal reduction and compaction using dissolution into acetone or water. The MgAl-CFX and ZrP-GTM fillers were obtained from the manufacturer Prolabin & Tefarm S.r.l. and their production has been previously described [22, 23]. The fillers had ∼50% w/w lamellar compounds (MgAl or ZrP) and the rest antibiotic (CFX or GTM). The compounding was done using a lab-scale Twin Screw Extruder at temperatures ranging between 140 and 150 °C, followed by die extrusion and pelletization, as described previously [22, 23]. Compounding was done by the supplier Nadir S.r.l. and the composites were provided in pellet form (Figure 1C, D).

### Rheological analysis – shear rate variation

Oscillation shear tests with 1% strain amplitude were conducted using a TA Discovery HR-1 rheometer. Stainless steel parallel plates with 25 mm of diameter were used. Tests were carried out in a nitrogen atmosphere, loading time was kept consistent, and melts were visually inspected at the end to avoid polymer degradation. Excess material was placed between the plates and the plates were brought together until the material flowed out across the perimeter. The overflown material was then scraped off and the tests were carried out after allowing the temperatures to stabilize. The inter-plate gap used was in the range 0.7 mm – 1 mm. Viscosity was measured at 210 °C for all materials at various angular velocities in the range 0.1 to 100 rad/s. The 210 °C temperature was chosen based on the fact that at this temperature all materials had a molten appearance. Measurements were also made at the extrusion temperatures of the highest filler loading material for each filler, the extrusion temperatures being empirically determined for a newly developed printhead [11]. These measurements were made in the angular velocity range 0.1 to 628 rad/s. In both cases, Cox-Merz transformation was applied to convert the complex viscosity vs. angular velocity plots to dynamic viscosity vs. shear rate plots and Carreau fit parameters were calculated using the TA Instruments TRIOS software to fit the obtained data.

### Mechanical testing of bulk materials under compression and tension

Mechanical tests for bulk materials were carried out using an Instron universal mechanical test machine and samples were prepared using ISO standards (ISO 527, Type 5, 4 mm thick samples for tensile tests and ISO 604, 10 mm x 10 mm x 4 mm samples for Compression tests). A 100 kN load cell was used and strain was applied at 0.1 mm/min (0.04 % strain/s). The samples were prepared using molds and a Babyplast micro-injection machine and working temperatures in the range 165-190 °C. The 15rGO composite could not be processed using the micro-injection machine due to its high viscosity and hence was not tested.

### Mechanical testing of scaffolds under compression

Scaffolds were manufactured with an overall dimensions of 20×20×4 mm^3^ and composed of filament meanders deposited in a 0-90 pattern, i.e. each layer had parallel filaments and the alternate layers had filament orientation rotated by 90° with respect to the other. The filament diameters were determined by the extrusion needles used (internal diameter, ID = 250 μm was used for all conditions, except for the fillers with the antibiotics, where a higher diameter needle, ID = 340 μm, was used to be able to extrude at relatively lower temperatures to avoid potential antibiotic degradation). The center-to-center spacing between the filaments was 750 μm for all scaffolds, except for those with antibiotics, where it was 850 μm, keeping the pore sizes constant at ∼500 μm. The layer height was 250 μm for the antibiotics containing scaffolds and 200 μm for the rest, thereby pushing each layer slightly into the previous layer for better bonding. PEOT/PBT scaffolds were also printed with the antibiotic containing scaffolds settings for their comparison, i.e. needle ID = 340 μm, 850 μm filament spacing and 250 μm layer height. Cylindrical samples with 4 mm diameter were cored out from the scaffolds using a biopsy punch and used for the mechanical tests (Figure 1E-H).

The scaffolds made from the highest loading of each type of filler were mechanically tested under compression, since based on the bulk properties, those would give the scaffolds with the highest moduli. For each material, the mechanical test samples were cored out from a single printed scaffold. For rGO, 10rGO was used instead of 15rGO, since 15rGO scaffolds were hard to core out test samples from, due to low interlayer bonding strength. The tests were carried out using an Instron Universal Mechanical Test machine, equipped with a 100 N load cell and 0.1 mm/min (0.04 % strain/s) strain rate. For these scaffolds, strength was calculated at yield, determined as the first local stress maxima.

Since printing reproducibility can also affect scaffold properties, scaffold samples from separate print batches were also tested. In this test, intermediate filler loading compositions were also included, as were 15rGO scaffolds that retained full integrity, which was achievable for a small fraction of cored out samples. These mechanical tests were carried out using a TA ElectroForce 3200 mechanical tester using a 450 N or a 50 N load cell and 1% strain/s strain rates. Elastic modulus was calculated as the slope of the stress-strain curves between 2 and 4% strains, where all curves were linear. Strength was calculated at failure (determined as the first occurrence of the stress dipping by 1% or more compared to the nearest stress maxima). Strength at yield was not evaluated, as the yield point (local stress maxima) was found sensitive to the noise filter used, while the strength at failure with a 1% drop as the threshold was found to be a robust measure. Although, a failure point was not found on all stress-strain plots (since sometimes the scaffolds smoothly transitioned from load bearing as porous structures to collapsed pore, bulk-like load bearing).

### Thermal and rheological analysis – temperature variation

The rheological analysis was carried out using an Advanced Rheometric Extended System (ARES, TA Instruments, USA) rheometer, equipped with a convection oven for temperature control. The measurements were performed using a parallel plates geometry with plates of 8 mm diameter. A gap of 0.8 mm was chosen and was kept constant in the non-isothermal tests, through the automatic adjustment of the tool thermal expansion.

Preliminary strain sweep tests were carried out to choose amplitudes that guaranteed conditions of linear viscoelasticity. For each measurement, a fresh sample was used, and the absence of degradation phenomena was checked at the beginning and at the end of the test. Cooling / heating cycles were carried out at 10 °C/min. The rheological transition temperatures from solid to liquid and from liquid to solid states were determined graphically, extrapolating the points of tangency between a straight line and the sigmoidal curve, in the inflection zone of the complex modulus G*.

Differential Scanning Calorimetry (DSC) tests were performed using a Mettler Toledo DSC-822. Thermal cycles at 10 °C/min were performed. The first heating ramp, carried out to remove the thermal history of the material, was discarded. The cooling and the second heating ramps were analyzed. Also in this case the solid-to-liquid and liquid-to-solid transition temperatures were determined in the maximum and minimum points of the heat flux versus the temperature plots.

### Computational models to predict mechanical properties of multi-material scaffolds

For single material scaffolds, the bulk mechanical properties provide an estimate of the scaffold properties that can be expected and mechanically testing scaffolds additionally provides information about the added effects on scaffold mechanical properties of AM process dependent factors, such as AM manufacturing errors and bond strength between layers. For multi-material scaffolds, the geometrical distribution of materials is an additional factor affecting the final scaffold mechanical properties. While the AM process effects on mechanical properties in going from bulk material to scaffolds cannot always be predicted, geometrical distribution of materials in scaffolds is straightforward and hence it should be possible to use computational modeling to predict multi-material scaffold properties from single material bulk and scaffold mechanical properties that were measured. This was tested using simplified geometry models where the scaffold porous architecture was replaced by bulk materials with measured scaffold mechanical properties and occupying the same total space as the given material and pores within the scaffolds.

The multi-material scaffold used was a 4 mm diameter, 15 mm high cylinder with three zones along the scaffold axis – the central zone was made of either PEOT/PBT or 45nHA and the two ends were made of the other material. The scaffolds were produced using a recently developed multi-material print head that could combine or switch continuously between two materials during a FDM AM process [11].

The model was divided into four types of material zones (Figure 6) – two zones for each of the materials and within each material zone, a cortical zone and a central zone with the properties of scaffold and bulk material respectively. The cortical region was given bulk material properties as it had low porosity and large overlaps between filaments of consecutive layers. This resulted from a modified print path in the cortical region, deviating from the 0-90 pattern elsewhere, in order to create supports for the next layer, as these scaffolds were individually printed and not punched out. The central zone had 0-90 pattern scaffolds of either material with the same architecture as the single material scaffolds tested, i.e. 250 μm diameter filaments, 750 μm inter-filament spacing, and 200 μm layer height. Computational models were solved using ANSYS Structural Mechanics. SOLID187 tetrahedral quadratic elements were used to account for the complex 3D shape changes.

## Results and Discussion

### Enhanced filler loading using plasticizer

The highest HA filler loadings possible using normal solvent blending, such that the final composite had sufficient integrity for processing and mechanical testing, were 45% HA w/w for mHA and 55% HA w/w for nHA. The use of the plasticizer allowed highest loading to reach 65% nHA w/w. HA micro-particles absorbed the liquid plasticizers, so they were not used. In addition, most of the plasticizers did not mix well with the polymer even with melt compounding, so triethyl citrate (TEC) was finally selected as the best performer. The finally selected composition for mechanical testing was 17.5% w/w each of PEOT/PBT and the TEC plasticizer and 65% w/w of nHA. In comparison, the highest HA loading that has previously been achieved in a thermoplastic polymer, where melt extrusion was also demonstrated, was 50% w/w with poly(ε-caprolactone) (PCL) [28]. For the mechanical characterization in this study, 45% w/w HA was selected as the highest loading for both mHA and nHA without plasticizer, to be able to compare the effect of particle size on mechanical properties. 65% w/w nHA with the plasticizer was the overall highest loading fraction tested for mechanical properties.

### Rheological analysis – shear rate variation

The reported composites were developed to be used with a screw extrusion based AM equipment, where the screw had regions with various shear rates [11]. To model computationally the extrusion process in the screw and to obtain reliable measures of operational torques for the motor, it was important to know the viscosity of the materials at various shear rates [11]. A straightforward way to implement the measured viscosity variations with respect to the shear rates in the models was to fit the viscosity vs. shear rate curves to the Carreau model (Equation 1) and provide the fit parameters in the fluid dynamics models, solved using COMSOL [11]. Thus, the Carreau parameters were calculated for all the materials (Supplementary Tables S2 and S3). This model fits the data to an equation (Equation 1) that predicts constant viscosities at very low and very high shear rates (Newtonian fluid), with a transition to an exponential change (power law fluid) for intermediate shear rates.

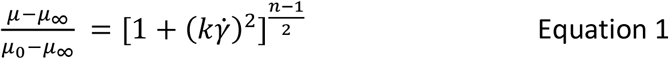

where *μ* is the viscosity, 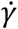 is the shear rate, *μ*_0_is the zero shear rate viscosity, *μ*_∞_ is the infinite shear rate viscosity, the term k is known as consistency, and n is the rate index. Where measurements were repeated, the fitted model parameters showed that while the viscosities at very high and very low shear rates, as predicted by the model had similar values in replicates, the transition positions and rates were not always consistent between measures. While the behavior of most materials fitted the Carreau models very well (R^2^>0.9), the polymer did not, and since it had low variations in viscosity with respect to shear rate, it would be better modelled as a constant viscosity material. It is to be noted here that the Cox-Merz transformation, which was used to obtain the Carreau fits, is generally found to not be applicable to high filler loading composites [29]. For high filler loading composites, the Cox-Merz transformation slightly overestimates viscosities [29]. Since this would only lead to slightly conservative estimates of extrusion screw parameters, it was acceptable to be used here nonetheless.

In general, the viscosity measurements of the materials with shear rate variations showed that the filler-loaded materials had a loading fraction proportional increase in viscosity compared to the polymer, but became less viscous at higher shear rates (shear-thinning) (Figure 2A-D, Supplementary figure S1). This provided with confidence for the empirical AM extrusion testing for the high rGO and HA filler loadings. For these fillers, the viscosities at low shear rates went up sharply with increasing filler loadings. In the high shear rate regions in the screw chamber, high viscosities challenge flow, but shear thinning is expected to assist.

**Figure 2.**
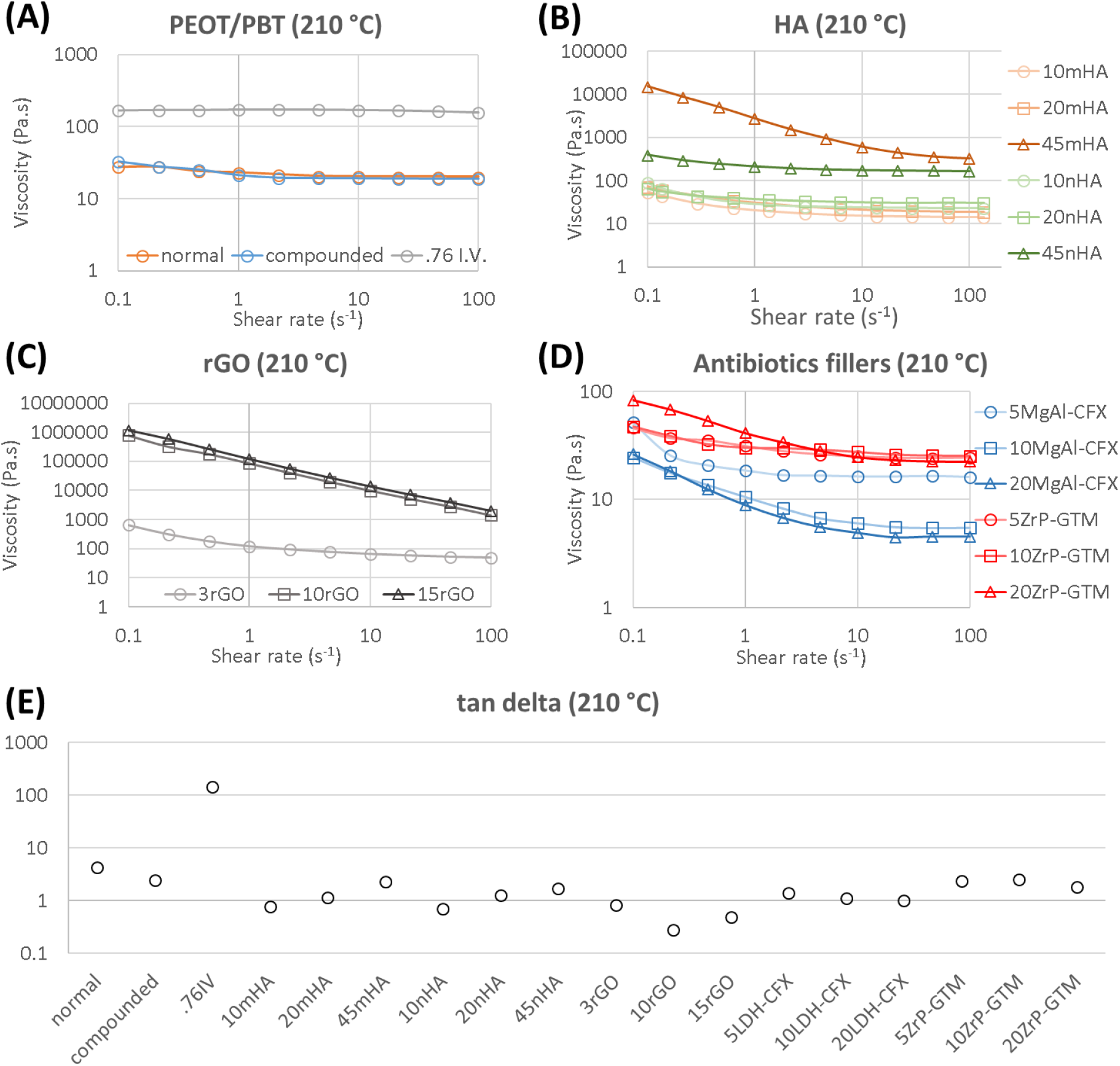
Viscosity (Pa.s) variations with shear rate (s^-1^), obtained by applying the Cox-Merz transform to the frequency sweep data of an oscillatory shear test, measured at 210 °C are shown for (A) PEOT/PBT, (B) mHA and nHA composites, (C) rGO composites and (D) MgAl-CFX and ZrP-GTM composites, for the various filler fractions tested. The tan delta values at the lowest shear rate (0.1 s^-1^) are compared (E) as measures of the extruded polymer melts behaving more like liquids (tan delta >1) or solids (tan delta < 1). The .76IV refers to the higher intrinsic viscosity PEOT/PBT (0.76 dl/g, vs. 0.51 dl/g for normal) tested for potential improvement in mechanical properties.

The results agreed with previously reported rheological studies of highly filler-loaded polymer composite melts [30], as well as more specifically the melts of HA loaded thermoplastic polymers, such as PCL [30-32]. These studies also found that filler addition increased the viscosity in proportion to filler volume fraction and this increase became sharper beyond ∼20% volume fraction (∼40% weight fraction for HA) [30, 32], and all the composites displayed shear thinning behavior, similar to the observations of this study. In addition, previous studies predicted higher viscosities for nanoparticle loadings than the same weight fraction of microparticles [30, 33], but in this study the opposite trend was observed for the HA composites at the highest loading fraction. The proposed explanation for the trend in literature was the higher surface area of the nanoparticles increasing the solid-solid friction. Along the same lines, the observation in this study could be explained by the greater particle aggregation observed in 45nHA than in 45mHA [34], thereby reducing the surface area for solid-solid friction. Due to the particle size and lamellar structure of rGO giving it a much higher surface area than the other fillers, the viscosity increase of rGO composites appeared at lower volume fractions. This is in agreement with previous reports on the rheology of polymer composites with graphene based materials [35] and also reflected by the bulk density of rGO being an order of magnitude smaller than the actual density, showing that a small mass of material can fill a large volume while being interconnected.

In addition, we also showed that, with a few exceptions, all materials had loss moduli (G”) consistently larger than the storage moduli (G’), i.e. tan delta (G”/G’) was generally greater than 1, showing that the composite melts generally behaved more as viscous liquids than solids (Figure 2E, Supplementary figures S2 and S3). The only exceptions were the composites 10mHA, 10rGO or 15rGO, where G”<G’ or tan delta was < 1. This effect also became apparent during the scaffold manufacture, mainly for the 15rGO composite. The consecutive layers did not bind well to each other, probably because of the poor mixing between filaments in alternate layers before cooling, as a result of the more solid like rheological behavior. This issue was not observed for the other materials, possibly because they were softer (lower storage modulus) than the 15rGO and had sufficient tack as per the Dahlquist criterion [24] for pressure sensitive adhesives, which states that materials with G’<0.1MPa have good adhesion. With shear rate variation, the tan delta increased at high shear rates in general, confirming the shear thinning behavior of the composite melts. It is to be noted here that the results for the highest fraction loading for each filler at the empirically determined extrusion temperatures have been reported previously [11], but are included here for completeness and comparison.

### Mechanical testing of bulk materials under compression and tension

The mechanical tests showed that increasing amounts of HA and rGO fillers generally made the composite stiffer than the bare polymer, but also more brittle compared to the polymer, thus failing at lower strains (Figure 3C-I). These trends were consistent between the tests performed under compression (Figure 3C-E) and tension (Figure 3F-I).

**Figure 3.**
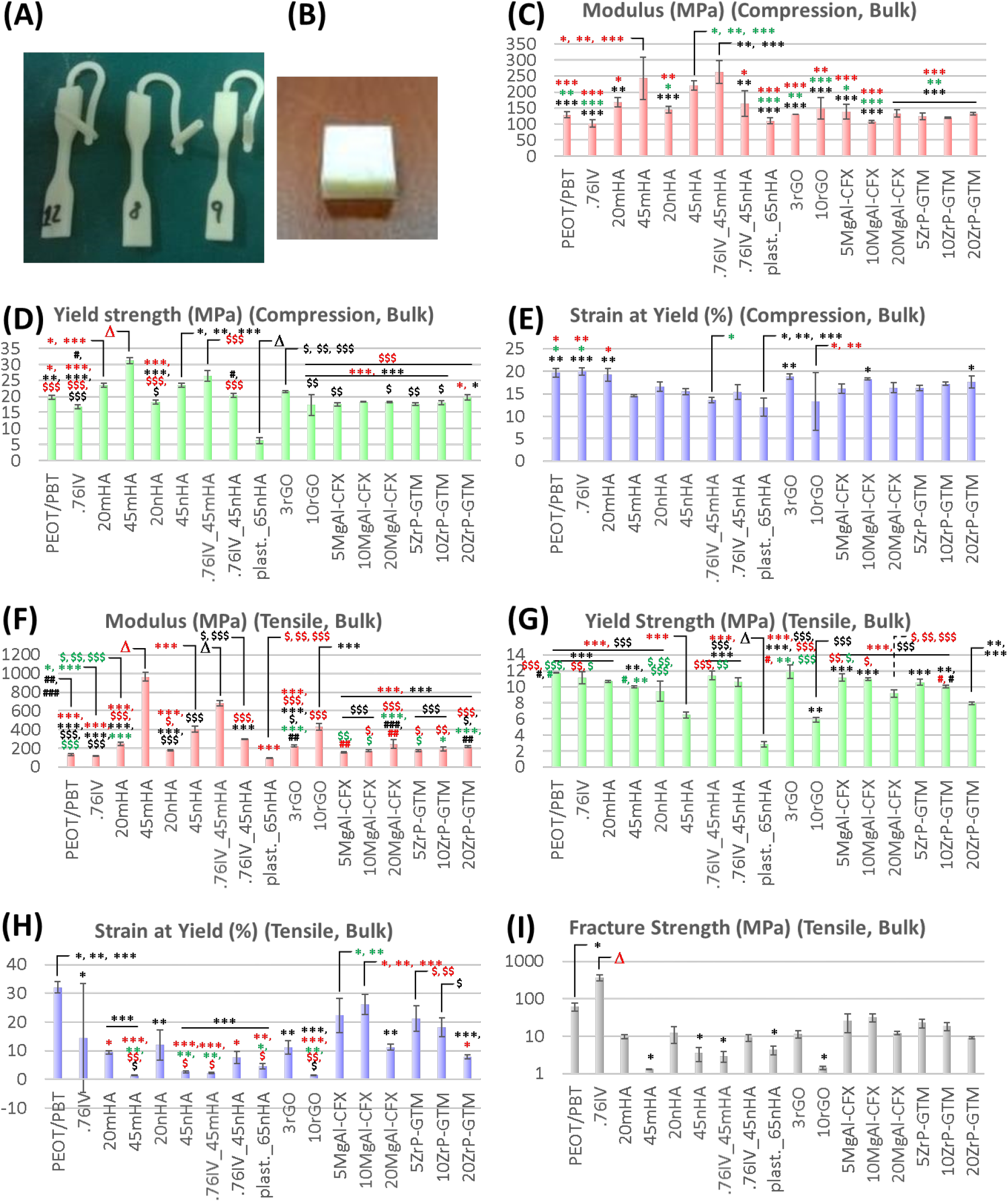
Bulk mechanical test samples are shown – (A) tensile test samples and (B) compression test samples. The measured elastic moduli (C, F), yield strength (D, G), strain at yield (E, H) are reported for both compression (C-E) and tensile tests (F-H), and fracture strength (I) for only the tensile tests. ANOVA p-values were <0.05 for all plots. Significant differences found in a Tukey’s post-hoc test are reported (*/$/# denoting 0.01<p <0.05, **/$$/## denoting 0.001<p <0.01, ***/$$$/### denoting p <0.001 and Δ denoting p<0.001 compared to all other materials).

The higher molecular weight, but otherwise identical, polymer (I.V. 0.76 dl/g PEOT/PBT) showed higher failure strength under tension than the default polymer (Figure 3I), and was hence tried as a means to improve the overall toughness of the 45% HA composites, which were the stiffest among those tested. However, this provided a statistically significant, yet only a marginal increase in the yield strength, only in tension and only for the 45nHA composite (Figure 3I).

Lastly, the composites generally showed higher load bearing ability but lower deformability in tension than in compression, as shown by the higher moduli and strengths and lower yield strains in tension than in compression (Figure 3C-I). This was most noticeable for the 45mHA micro-particles composite.

The best performing materials in terms of increased load bearing ability were the 45% HA composites with a ∼1.7x increase over the polymer alone and comparable results for both particle sizes, in terms of the compressive modulus. The compressive moduli of these composites (220 ± 14 MPa for 45nHA and 242.5 ± 66 MPa for 45mHA) were in the range of trabecular bone compressive modulus (0.1 – 2 GPa). The results also compare well to previously reported ∼1.7x increase (498.3 MPa vs. 299.3 MPa) in modulus over polymer alone, for a loading of 30% HA (w/w) filler in PCL, which is a commonly used thermoplastic polymer for producing bone TE scaffolds [36].

### Mechanical testing of scaffolds under compression

Scaffolds were only tested under compression, since this is the primary mode of loading that they are subjected upon implantation in a skeletal location. As it is known that different production batches could lead to experimental variations, mechanical tests have been done on different scaffold batches. Samples from the same additive manufacturing batch showed low variability in the measured values of the mechanical properties (figure 4), while scaffolds taken from various printing batches showed larger variations in the measured values (supplementary figure S4). Amongst the highest filler loading scaffolds (from the same printing batches), the general trend of reduced yield strain of composites over polymer were consistent with the bulk materials (figure 4). Unlike for bulk materials, the moduli showed roughly similar values, as also observed previously for scaffolds produced from PCL and PCL-HA composites [37]. 45nHA and 20ZrP-GTM showed the opposite trend to bulk materials in terms of moduli, with significantly lower and higher moduli than polymer scaffolds, respectively. Since punching and handling errors could be ruled out due to the low standard deviations in the measurements, all these effects were attributed to precision errors in printing and material-specific effects such as bonding between printed filaments. The latter was further investigated by means of combined thermal-rheological analysis of nHA and ZrP-GTM composites.

**Figure 4.**
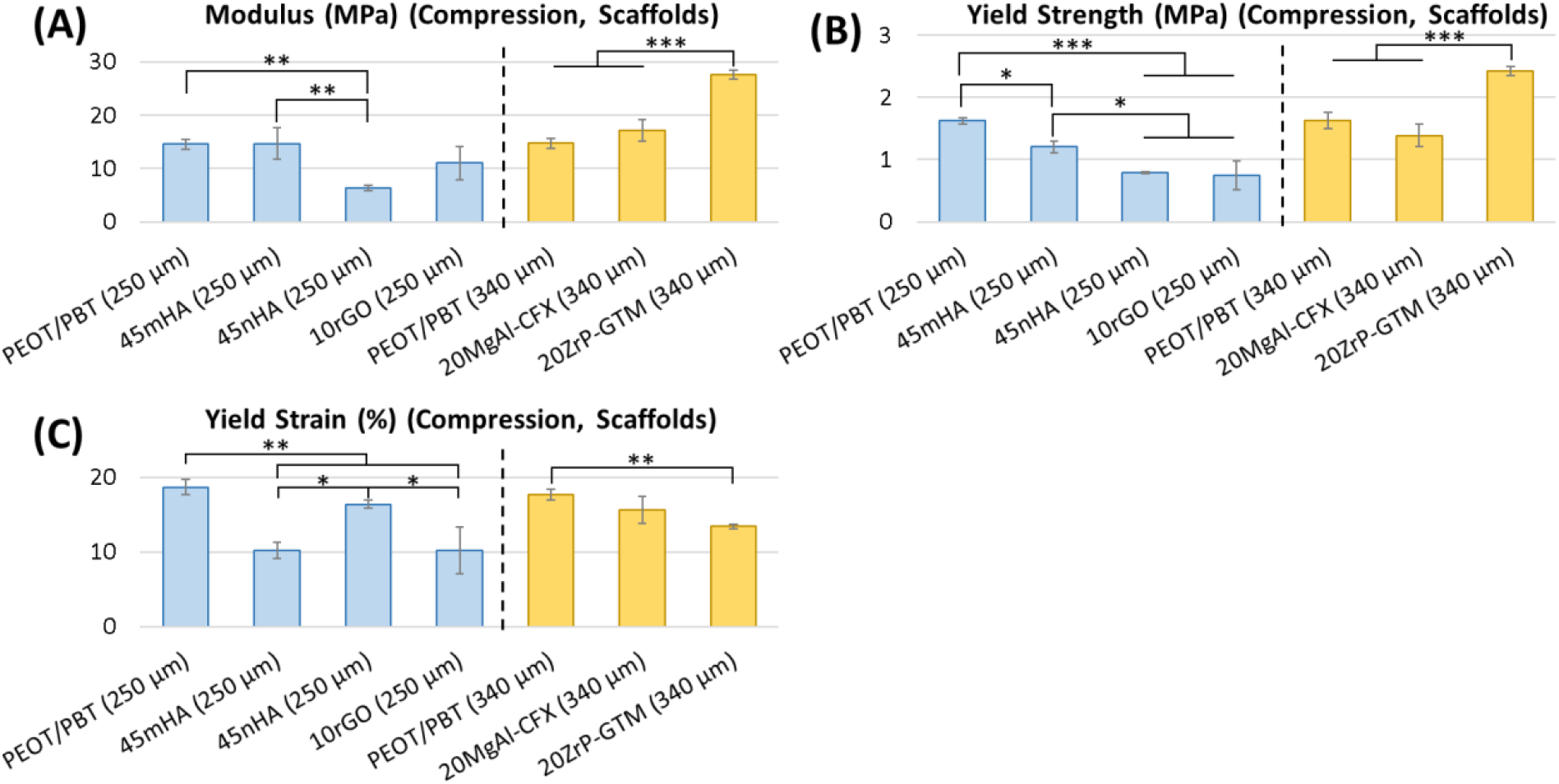
The elastic moduli (A), yield strength (B) and strain at yield (C) are reported for the high loadings scaffolds made of the various materials tested under compression. The results are separated for the scaffolds made using the two filament diameters – 250 μm and 340 μm. (**\***: p<0.05, **\*\***: p<0.01, **\*\*\***: p<0.001).

In the test taking into account the printing batch variability and intermediate filler concentrations, no significant differences were found between the moduli (Supplementary figure S4). Statistical analysis was not carried out for the failure strength and strain, since not all samples provided these measurements. In some samples, the load never dropped with increasing compression strain, as they smoothly transferred from pore collapse to load bearing as bulk materials. Nonetheless, two materials stood out with distinct properties – 3rGO scaffolds showed the highest failure strains (46.4 ± 9%) and 15rGO scaffolds showed the highest individual (53.1 MPa) and mean (33.9 ± 17.4 MPa) modulus among all samples. While the scaffold moduli (∼20 MPa) were below the trabecular bone properties, they could be brought back within the trabecular bone moduli range by lowering the porosity. More interestingly, we aim to investigate in future studies the improvements that can be brought by increasing the fiber overlap, without lowering porosity. This has been recently shown to be achievable by printing in hexagonal patterns [38].

### Thermal and rheological analysis – temperature variation

This analysis was conducted on the nHA and ZrP-GTM composites, mainly to evaluate if crystallization dynamics could explain why the improved bulk mechanical properties of nHA materials did not translate to scaffolds and why ZrP-GTM scaffolds appeared stiffer than the polymer when the bulk material did not show such an effect.

The rheological measurements with temperature ramps showed that for both materials, the addition of fillers led to the composite melts solidifying at higher temperatures than the polymer alone, as observed by filler-fraction proportional shifts in the sharp increases in viscosity during cooling ramps (Figure 5A, B, Table 1). This could be due to the filler assisted change in polymer crystallization, as it is known to occur due to filler particles acting as effective nucleation sites for polymer crystals to form and grow [39, 40]. DSC analysis supported this reasoning for both composites, since both of them showed an increase in the solidification temperature (Figure 5C, D, Table 2). Interestingly for nHA, the solidification temperature shift from that of the base polymer was much higher when predicted by rheology than by DSC, suggesting that nHA promotes (amorphous) solidification more strongly than ZrP-GTM, even if the effect on polymer crystallization was similar for the two fillers.

**Figure 5.**
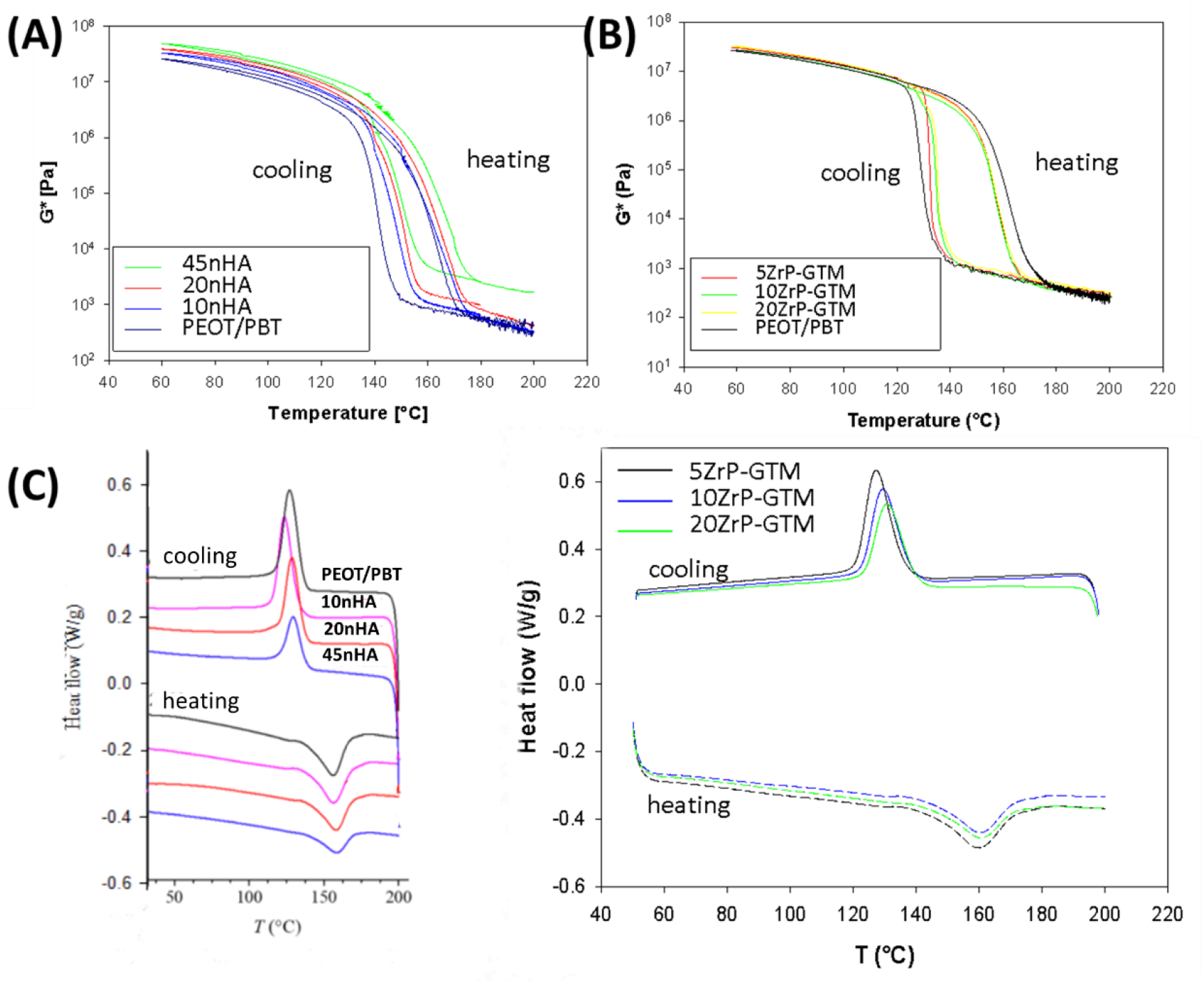
The variation of the complex moduli of (A) nHA and (B) ZrP-GTM composites is shown in heating-cooling cycles during the rheological measurements. DSC plots of the (C) nHA and (D) ZrP-GTM composites provide insights into the links between the shifts observed in the rheological measurements on filler addition and the polymer crystallization in the presence of the fillers.

**Table 1.** Liquid to solid (L->S) and solid to liquid (S->L) transition temperatures for the nHA and ZrP-GTM composites calculated from the rheological measurements

| **Sample** | **$T_{L \rightarrow S}$** | **$T_{S \rightarrow L}$** |
| --- | --- | --- |
| **PEOT/PBT** | 130 | 157 |
| **10nHA** | 141 | 159 |
| **20nHA** | 145 | 164 |
| **45nHA** | 143 | 162 |
| **5ZrP-GTM** | 133 | 156 |
| **10ZrP-GTM** | 135 | 158 |
| **20ZrP-GTM** | 136 | 159 |

**Table 2.** Liquid to solid (L->S) and solid to liquid (S->L) transition temperatures for the nHA and ZrP-GTM composites calculated from the DSC measurements

| **Sample** | **$T_{L \rightarrow S}$** | **$T_{S \rightarrow L}$** |
| --- | --- | --- |
| **PEOT/PBT** | 127 | 156 |
| **10nHA** | 123 | 156 |
| **20nHA** | 128 | 158 |
| **45nHA** | 129 | 159 |
| **5ZrP-GTM** | 127 | 160 |
| **10ZrP-GTM** | 129 | 160 |
| **20ZrP-GTM** | 131 | 161 |

The specific enthalpies calculated from the DSC measurements (Table 3) showed a decrease with increasing amounts of ZrP-GTM, suggesting a lower fraction of crystallized polymer or an effect of the lower mass fraction of the polymer. However, increasing fractions of nHA showed an increase in specific enthalpies of crystallization, suggesting a higher crystalline fraction. The faster solidification of nHA composites suggests that the binding between filaments of scaffolds might be weaker, leading to weaker scaffolds, but the higher crystallization of the polymer suggests that the scaffolds should have been stiffer than those without the nHA. The observed faster solidification and lower polymer crystallization for the ZrP-GTM composites suggests that the scaffolds should have been less stiff than without the filler. Thus, a change in polymer solidification and crystallization rates did not explain the observed anomalous results for the ZrP-GTM and nHA scaffolds. The thermal-rheological analysis should have picked up any major mechanical property changes in going from the bulk material to the scaffold, since they include both manipulations applied to the composites in the printing process, i.e. shear stress and temperature changes. Given that the anomalous results for nHA and ZrP-GTM do not appear in the mechanical investigation of scaffolds from multiple batches, the results of the single batch measurements could best be attributed to scaffold production process variability, leading to dimensional errors. Future studies should additionally include direct mechanical tests of extruded filaments and delamination tests to compare the bonding strengths between the printed layers. Micro computed tomography assisted actual geometry determination could also help control for effects of dimensional errors during printing. While not relevant for explaining the scaffold mechanical properties, the thermal-rheological analysis additionally showed that the addition of the fillers also increased the melting temperatures of the composites in proportion to the filler fraction and these shifts were similar for both materials using the two types of measurements (Tables 1 and 2).

**Table 3.** Enthalpy values extrapolated from DSC tests

| **Sample** | **$\Delta H$ ( J/g) Cooling** | **$\Delta H$ ( J/g) Melting** |
| --- | --- | --- |
| **0%** | 21.5 | -21.5 |
| **10% nHA** | 21.8 | -20.6 |
| **20% nHA** | 21.9 | -19.5 |
| **45% nHA** | 23 | -22 |
| **5% ZrP+GTM** | 19.3 | -17.8 |
| **10% ZrP+GTM** | 16.8 | -15.9 |
| **20% ZrP+GTM** | 16.4 | -14.3 |

### Simplified computational models to predict mechanical properties of multi-material scaffolds

The reported material library was developed also with an aim for producing multi-material scaffolds using a newly developed printhead [11]. It was therefore desirable to have a computational predictability of mechanical properties of such scaffolds in order to design multi-material scaffolds with improved mechanical properties without time consuming empirical optimization. Since scaffolds showed material-specific processing effects on mechanical properties, scaffold properties were used as inputs for models instead of bulk material properties, which also allowed for greatly simplifying the geometry. This was done for multi-material scaffolds made of alternating regions of the two materials – 45nHA composite or the PEOT/PBT polymer alone (figure 6A, B). The scaffolds were printed in the shape necessary to fill a long bone segmental defects in a rabbit *in vivo* model (4 mm diameter, 15 mm long cylinders). These scaffolds had peripheral filament regions with higher overlap between consecutive layers than central regions, in order to improve the stability of the scaffolds with otherwise low amount of load-bearing filament intersections, as a result of the scaffolds’ small size (figure 6C). Modeling this low porosity peripheral area with the bulk mechanical properties of the respective materials and the rest with the scaffold mechanical properties (figure 6D) gave a good prediction of the multi-material scaffold modulus. These results were closer to the measured multi-material scaffolds’ moduli than the full-geometry models using bulk material properties, irrespective of whether scaffolds’ moduli used were from the single-batch test or multi-batch test (Table 4). Furthermore, modeling the scaffolds as continuous materials with the measured scaffold mechanical properties was also computationally economical compared to modeling for the full filament structures of scaffolds. Solving for ∼2.7 million elements, which takes on the order of 30 minutes for linear analysis and weeks for a non-linear analysis, was thus reduced to a problem solvable with ∼10,000 elements, which require a few seconds to compute for a linear analysis and a few minutes to compute for a non-linear analysis (based on a 2.4 GHz processor and a 64 GB RAM system).

**Figure 6.**
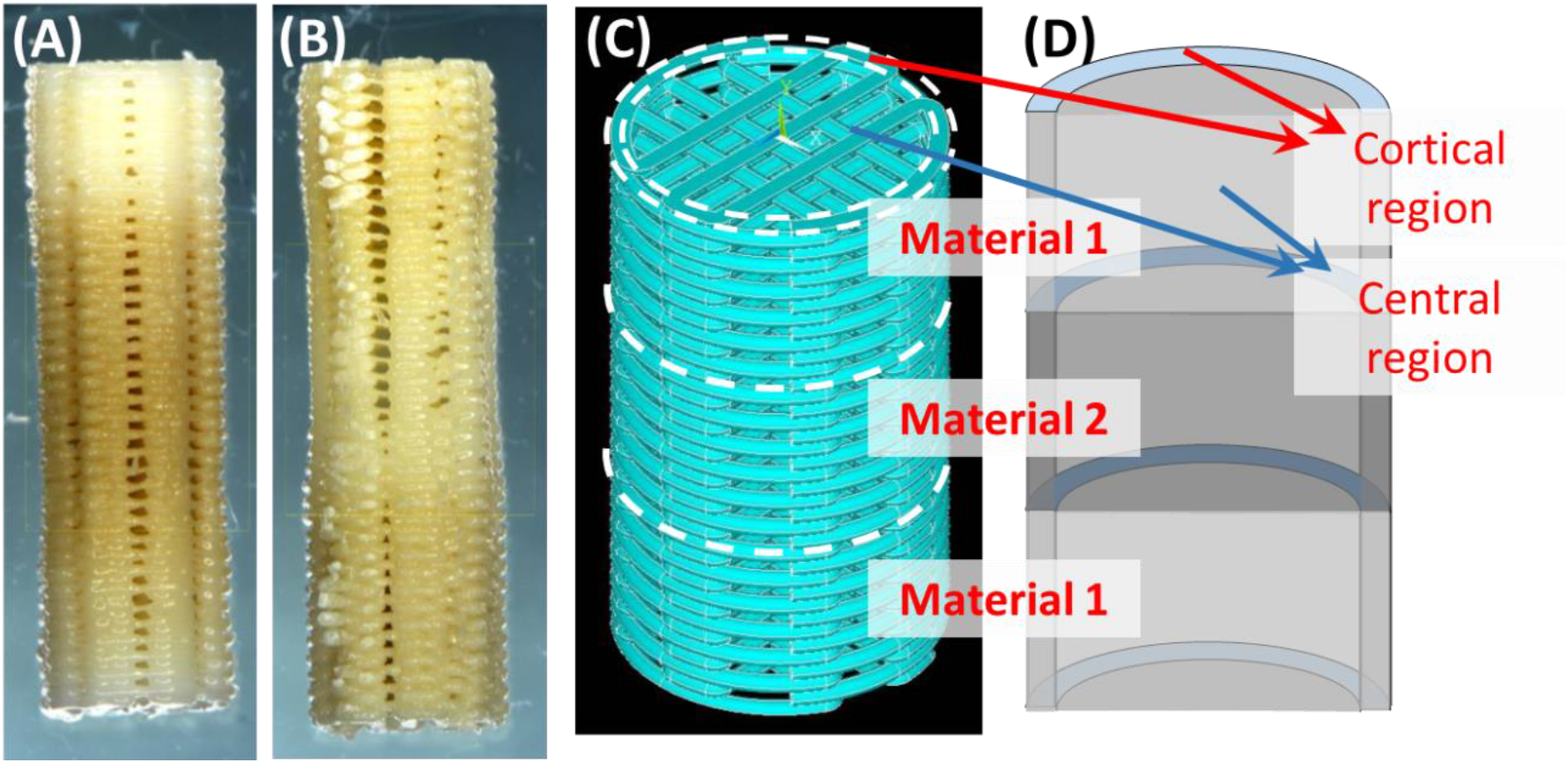
The multi-material scaffold with (A) 45nHA in the middle and (B) PEOT/PBT in the middle are shown. The scaffolds were mechanically tested and the results were compared with modeling results using measured bulk and scaffold mechanical properties. Models were solved using the full scaffold structure (C) or a simplified structure (D).

**Table 4.**
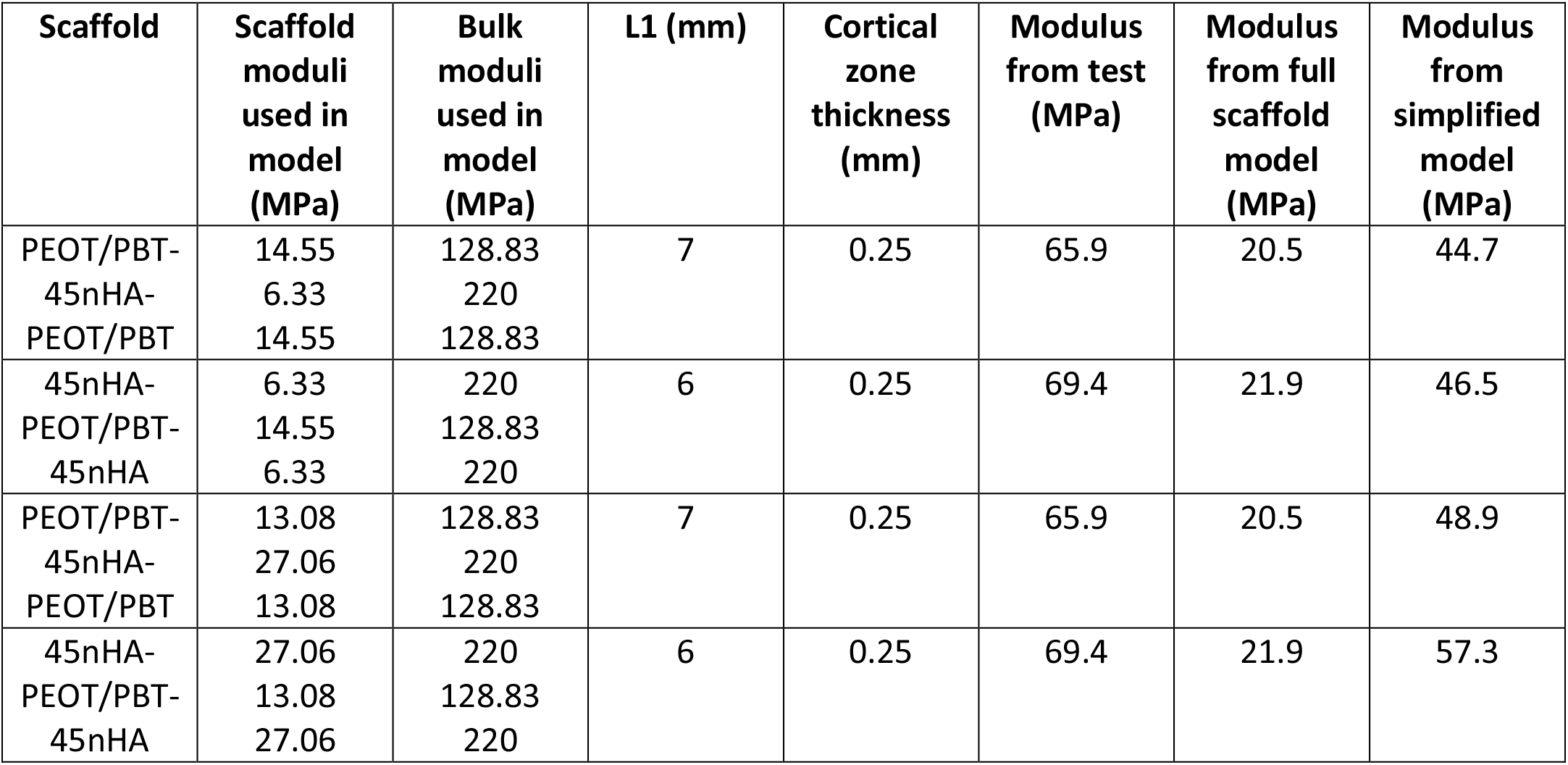
Model parameters and results

## Conclusion

We report here the rheological and mechanical characterization of a range of composite materials, formed by loading various fillers into a thermoplastic polymer, and developed for producing additively manufactured bone tissue engineering scaffolds. These characterizations are important for determining processability of the materials by the additive manufacturing technique of choice and give an estimate of the mechanical properties of the resulting scaffolds. The mechanical properties of rabbit long bone critical size defect scaffolds were also measured. The results show that increasing amounts of fillers sharply increase the viscosity of the composite material melts at high filler loadings. However, the materials show shear-thinning behavior, which allows for their processing using AM techniques that can apply high shear rates. The tan delta values (>1), showing more solid-like behavior of the 15rGO material melt, coupled with its high storage modulus, was predictive of the observed difficulty in bonding between 15rGO scaffold layers. In terms of mechanical properties, the fillers and concentrations tested could nearly double the modulus of the polymer without fillers, as observed for the 45% HA scaffolds under compression. However, between material-specific scaffold production process effects and batch-to-batch variability in scaffolds, the mechanical improvement seen in bulk materials seems to get masked in scaffolds. Combined thermal-rheological analysis of nHA and ZrP-GTM composites could not resolve the observation of anomalous scaffold mechanical properties, but still provided useful insights. Lastly, we show the utility of measuring both bulk and scaffold mechanical properties in predicting the mechanical properties of complex multi-material scaffolds using simple computational models. Overall, the results reported here will serve as an important guide to future researchers using the developed materials.

## Acknowledgements

The work was supported by a Horizon 2020 research and innovation programme grant from the European Union, called the FAST project (grant no. 685825, project website: http://project-fast.eu). The authors acknowledge the support of the FAST project consortium for the various aspects of this work.

## Author contributions

R.S. prepared the manuscript, carried out the rheological measurements with shear rate variations, produced scaffolds, and carried out scaffold mechanical tests. A.S. and S.V. prepared the various HA materials and carried out all other mechanical tests reported. M.C.-T. optimized printing parameters and printed the scaffolds tested. I.C.U.-A., A.M. and S.P. carried out the computational modeling. A.C. and J.H. designed protocols and carried out initial tests for the shear rate variation rheological study. A.G. and L.C. carried out the thermal and temperature variation rheological analysis on the antibiotics and HA materials respectively, under the supervision of V.V. and N.G.. M.Si. provided the antibiotics based fillers, R.W. provided the rGO material, and M.Sc. produced the antibiotics-and rGO-based composites using melt compounding. A.P., C.M. and L.M. supervised R.S., M.C.-T., and A.C., and designed the overall study for which the reported materials were developed.

