## Supplementary Material for "Additive manufactured scaffolds for bone tissue engineering: physical characterization of thermoplastic composites with functional fillers"

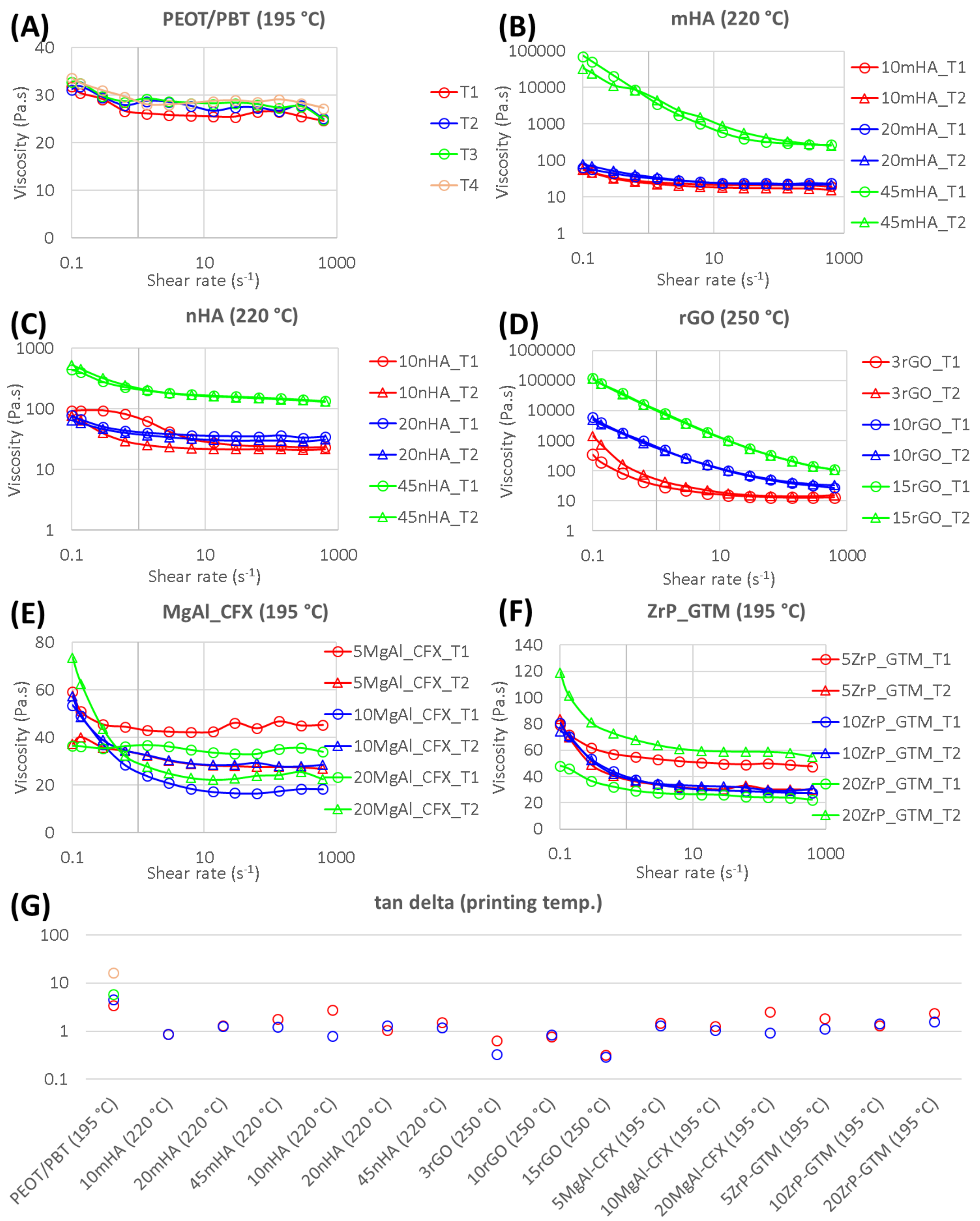


Figure S1 – Complex viscosity (Pa.s) variations with shear rate ($s^{-1}$), measures at empirically determined AM extrusion temperatures are shown for (A) PEOT/PBT, and (B) mHA, (C) nHA, (D) rGO, (E) MgAl-CFX, and (F) ZrP-GTM composites, for the various filler fractions tested. The tan delta values at the lowest shear rate (0.1 $s^{-1}$) are compared (G).

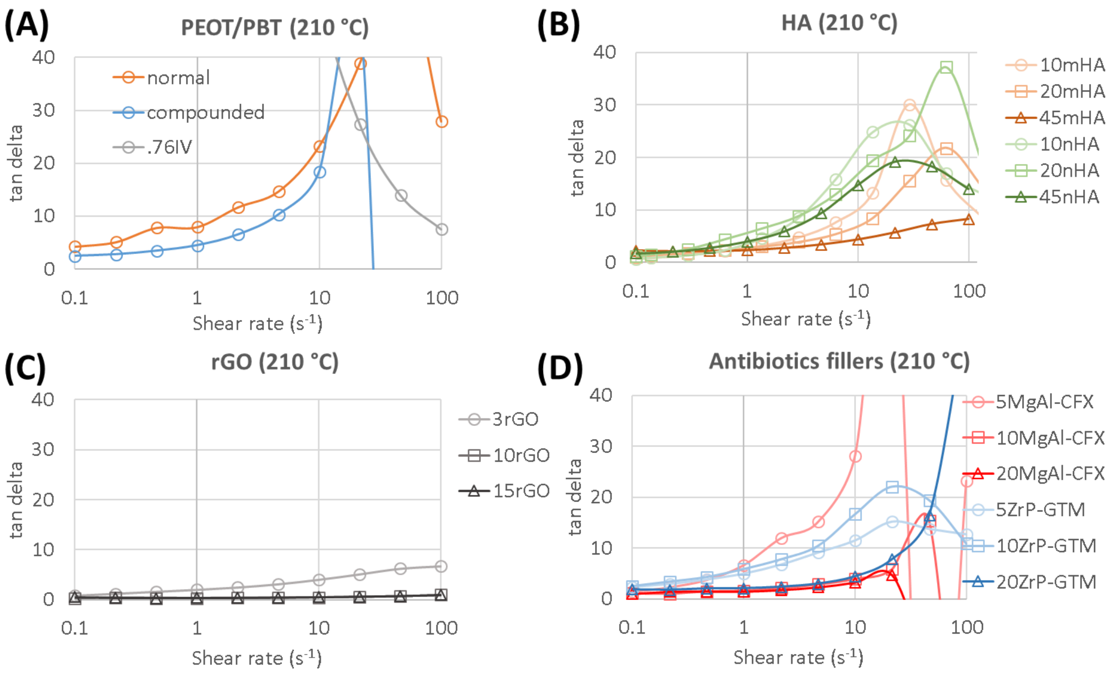


Figure S2 – Tan delta variations with shear rate ($s^{-1}$), at 210 °C, are shown for the various materials and compositions tested. Extremely high and negative values were recorded at high shear rates and for .76IV material at low shear rates. Those values were most likely a result of experimental noise and hence Y-axis scales are chosen ignoring those values, and rather to highlight trends in regions where the measurements were reliable.

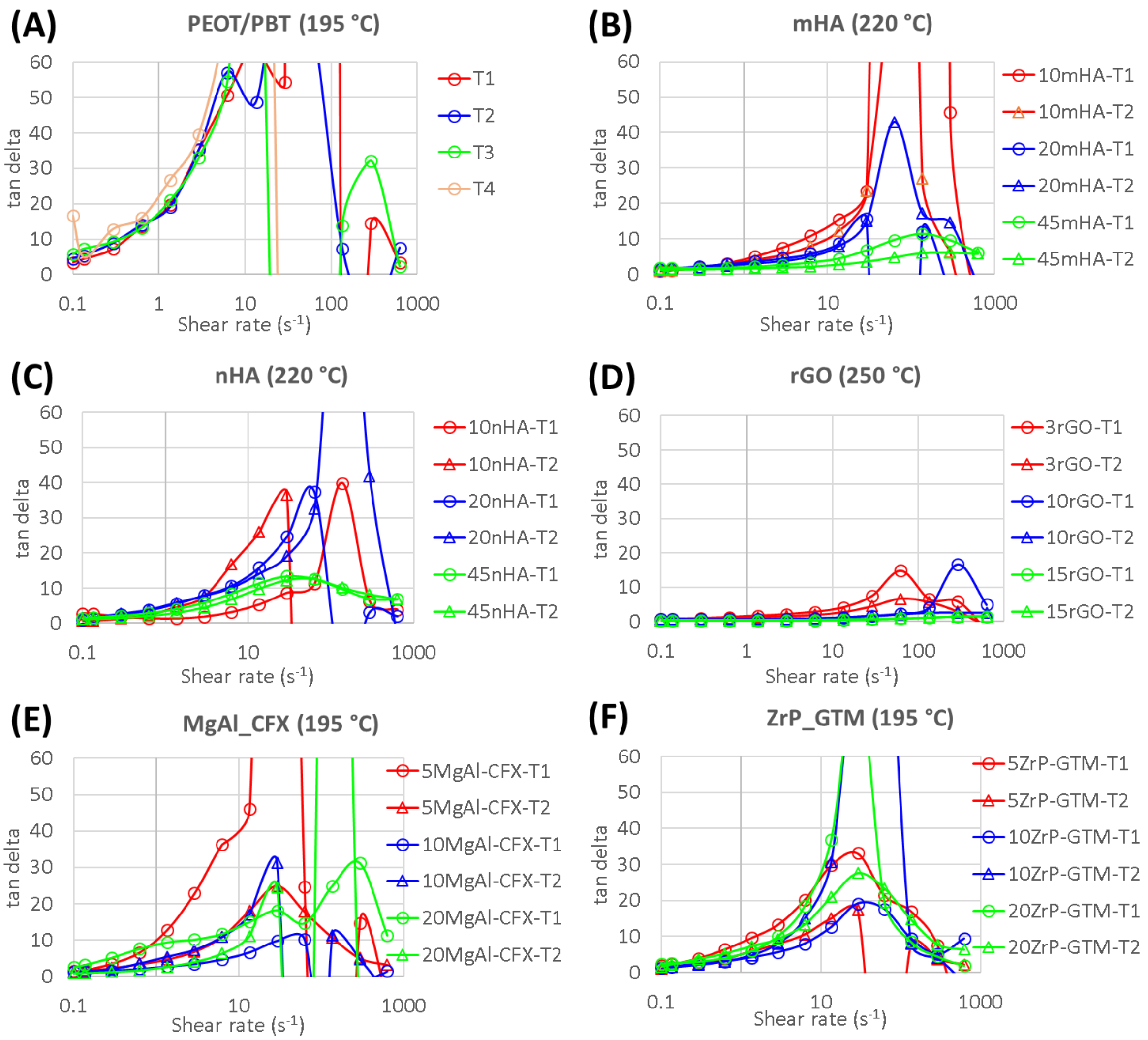


Figure S3 - Tan delta variations with shear rate ($s^{-1}$), at empirically determined AM extrusion temperatures, are shown for the various materials and compositions tested. Extremely high and negative values were again recorded at high shear rates. Those values being the ones where the plots did not overlap between repeats for the same material, confirm that they were a result of experimental noise. Hence, again, Y-axis scales are chosen ignoring those values, focusing on the trends in regions where the measurements were reliable.

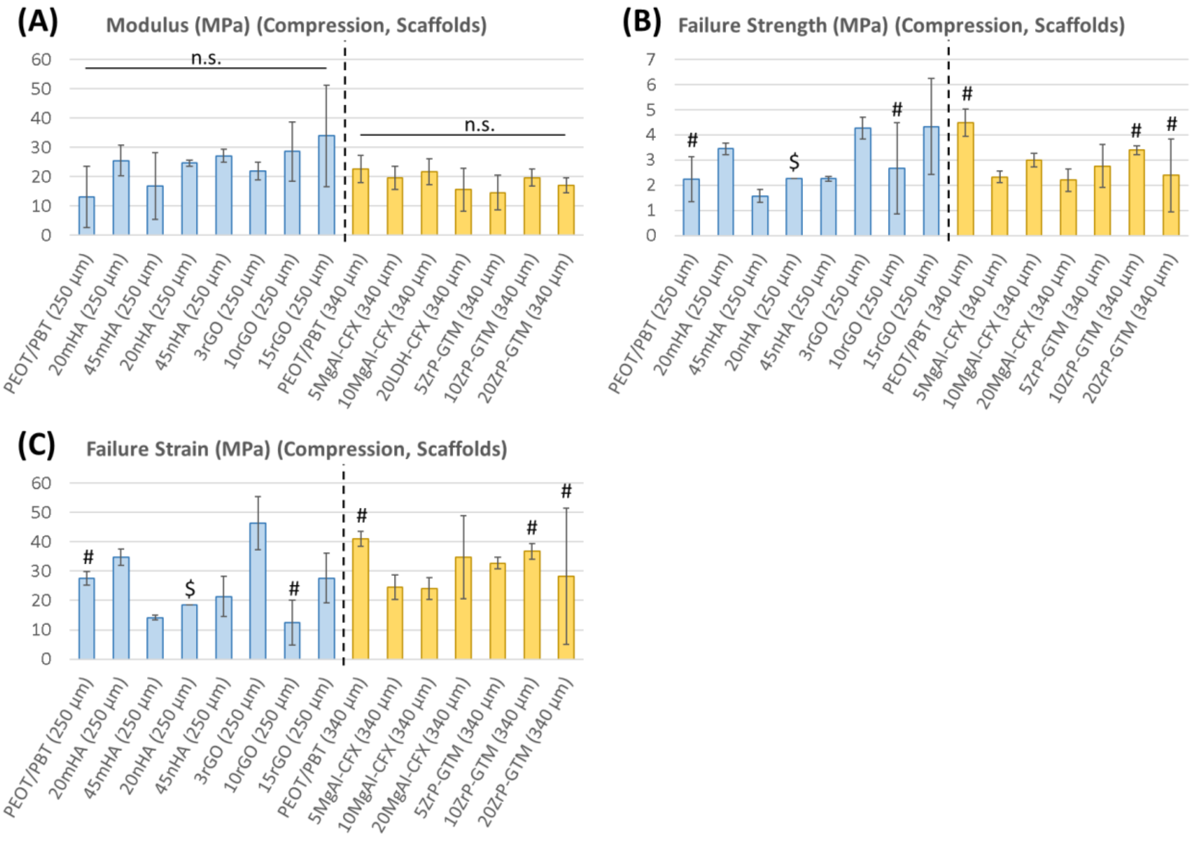


Figure S4 – The elastic moduli (A), failure strength (B) and strain at failure (C) are reported for scaffolds made of the various materials tested under compression. The results are separated for the scaffolds made using the two filament diameters - 250µm and 340 µm. (**n.s.**: not statistically significant, **#**: n=2, **$**: n=1).

Table S1 – Composite materials nomenclature

| Short name | Polymer / Composite material composition |
|---|---|
| **PEOT/PBT** or **normal** | poly(ethylene oxide terephthalate) / poly(butylene terephthalate block co-polymer prepared from 300 g/mol PEO and with PEOT:PBT w/w ratio 55:45, and molecular weight dependent intrinsic viscosity of 0.51 dl/g, unless specified |
| **compounded** | PEOT/PBT processed in twin screw compounder |
| **.76IV** | higher molecular weight PEOT/PBT (intrinsic viscosity 0.76 dl/g) |
| **10mHA, 20mHA, 45mHA** | 10, 20 and 45% (w/w) hydroxyapatite microparticles (5 $\pm$ 1 μm) composite with PEOT/PBT (I.V. 0.51 dl/g, unless specified) |
| **10nHA, 20nHA, 45nHA** | 10, 20 and 45% (w/w) hydroxyapatite nanoparticles (<200 nm) composite with PEOT/PBT (I.V. 0.51 dl/g, unless specified) |
| **Plast.-65nHA** | 65% (w/w) hydroxyapatite nanoparticles (<200 nm), and 17.5% (w/w) triethyl citrate (TEC) plasticizer composite with PEOT/PBT (I.V. 0.51 dl/g) |
| **5ZrP-GTM, 10ZrP-GTM, 20ZrP-GTM** | 5, 10 and 20% (w/w) gentamycin intercalated in zirconium phosphate composite with PEOT/PBT (I.V. 0.51 dl/g) |
| **5MgAl-CFX, 10MgAl-CFX, 20MgAl-CFX,** | 5, 10 and 20% (w/w) ciprofloxacin intercalated in MgAl layered double hydroxide composite with PEOT/PBT (I.V. 0.51 dl/g) |
| **3rGO, 10rGO, 15rGO** | 3, 10 and 15% (w/w) reduced graphene oxide composite with PEOT/PBT (I.V. 0.51 dl/g) |

Table S2 – Carreau parameters for the various materials at 210 °C.

| sample | shear rate range | temperature | Carreau parameters | | | | |
|---|---|---|---|---|---|---|---|
| | | | $\mu_0$ (Pa.s) | $\mu_\infty$ (Pa.s) | consistency (s) | rate index | $R^2$ |
| compounded-PEOT/PBT | 0.1 to 100 | 210 | 34.35 | 19.11 | 4.65 | -0.14 | 0.99 |
| PEOT/PBT | 0.1 to 100 | 210 | 28.65 | 20.05 | 3.6 | 0.17 | 0.98 |
| 0.76 IV | 0.1 to 100 | 210 | 170.25 | 178.84 | 3.64E-11 | 0.05 | -6.66E-16 |
| 45mHA | 0.1 to 100 | 210 | 24136.5 | 276.01 | 14.44 | 0.15 | 1 |
| 45nHA | 0.1 to 100 | 210 | 7355.41 | 168.78 | 1633.1 | 0.32 | 1 |
| 3rGO | 0.1 to 100 | 210 | 91500.9 | 51.77 | 3506.33 | 0.13 | 0.99 |
| 10rGO | 0.1 to 100 | 210 | 159000000 | 312.47 | 3178.77 | 0.06 | 1 |
| 15rGO | 0.1 to 100 | 210 | 181000000 | 675.75 | 1768.57 | 0.02 | 1 |
| 5-MgAl-CFX | 0.1 to 100 | 210 | 40293 | 16.54 | 1067.12 | -0.52 | 0.99 |
| 10-MgAl-CFX | 0.1 to 100 | 210 | 30.54 | 5.03 | 11.68 | 0.35 | 1 |
| 20-MgAl-CFX | 0.1 to 100 | 210 | 33.55 | 4.28 | 10.37 | 0.19 | 1 |
| 5-ZrP-GTM | 0.1 to 100 | 210 | 134.72 | 22.96 | 277.66 | 0.52 | 0.99 |
| 10-ZrP-GTM | 0.1 to 100 | 210 | 404.65 | 25.54 | 1177.79 | 0.39 | 0.98 |
| 20-ZrP-GTM | 0.1 to 100 | 210 | 89.67 | 20.95 | 6.56 | 0.35 | 1 |

Table S3 - Carreau parameters for the various materials at empirically determined AM extrusion temperatures.

| sample | shear rate range | temperature | Carreau parameters | | | | |
|---|---|---|---|---|---|---|---|
| | | | $\mu_0$ (Pa.s) | $\mu_\infty$ (Pa.s) | consistency (s) | rate index | $R^2$ |
| PEOT/PBT_T1 | 0.1 to 627.997 | 195 | 26.92 | 15.28 | 3.87E-08 | -0.08 | 3.83E-10 |
| PEOT/PBT_T2 | 0.1 to 627.997 | 195 | 45.62 | 26.32 | 555.8 | 0.67 | 0.8 |
| PEOT/PBT_T3 | 0.1 to 627.997 | 195 | 44.02 | 26.07 | 696.45 | 0.75 | 0.8 |
| PEOT/PBT_T4 | 0.1 to 627.997 | 195 | 33.98 | 28.46 | 3.2 | -1.27 | 0.92 |
| 10mHA_T1 | 0.1 to 627.997 | 220 | 3529.02 | 20.82 | 2312.21 | 0.16 | 0.99 |
| 10mHA_T2 | 0.1 to 627.997 | 220 | 1401.12 | 16.54 | 1399.4 | 0.26 | 0.99 |
| 20mHA_T1 | 0.1 to 627.997 | 220 | 81.96 | 23.2 | 15.22 | 0.34 | 1 |
| 20mHA_T2 | 0.1 to 627.997 | 220 | 108.39 | 21.61 | 17.68 | 0.38 | 1 |
| 45mHA_T1 | 0.1 to 627.997 | 220 | 29000000 | 273.58 | 2050.2 | -0.13 | 1 |
| 45mHA_T2 | 0.1 to 627.997 | 220 | 1545000 | 216.104 | 1461.63 | 0.22 | 1 |
| 10nHA_T1 | 0.1 to 627.997 | 220 | 95.64 | 23.66 | 1.05 | -0.13 | 1 |
| 10nHA_T2 | 0.1 to 627.997 | 220 | 146.34 | 21.64 | 18.07 | -0.12 | 1 |
| 20nHA_T1 | 0.1 to 627.997 | 220 | 3165.35 | 35.01 | 1700.46 | 0.16 | 0.99 |
| 20nHA_T2 | 0.1 to 627.997 | 220 | 526.7 | 29.88 | 632.92 | 0.36 | 1 |
| 45nHA_T1 | 0.1 to 627.997 | 220 | 5792.7 | 145.252 | 1151.8 | 0.38 | 0.99 |
| 45nHA_T2 | 0.1 to 627.997 | 220 | 9839.54 | 140.06 | 1311.61 | 0.34 | 1 |
| 3rGO_T1 | 0.1 to 627.997 | 250 | 131903 | 13.65 | 2245.46 | -0.14 | 0.99 |
| 3rGO_T2 | 0.1 to 627.997 | 250 | 1627430 | 15.79 | 1288.64 | -0.5 | 0.99 |
| 10rGO_T1 | 0.1 to 627.997 | 250 | 630497 | 28.65 | 2273.45 | 0.11 | 1 |
| 10rGO_T2 | 0.1 to 627.997 | 250 | 633633 | 33.03 | 3174.36 | 0.14 | 1 |
| 15rGO_T1 | 0.1 to 627.997 | 250 | 41500000 | 94.03 | 4946.38 | 0.04 | 1 |
| 15rGO_T2 | 0.1 to 627.997 | 250 | 41100000 | 93.72 | 5155.58 | 0.04 | 1 |
| 5MgAl-CFX_T1 | 0.1 to 627.997 | 195 | 85.8 | 44.28 | 1.68 | -72.24 | 0.87 |
| 5MgAl-CFX_T2 | 0.1 to 627.997 | 195 | 38.9 | 27 | 3.71 | 0.46 | 0.98 |

| | | | | | | | |
|---|---|---|---|---|---|---|---|
| 10MgAl-CFX_T1 | 0.1 to 627.997 | 195 | 61.01 | 17.25 | 7.62 | 0.14 | 0.99 |
| 10MgAl-CFX_T2 | 0.1 to 627.997 | 195 | 1567.43 | 28.43 | 1357.4 | 0.18 | 0.99 |
| 20MgAl-CFX_T1 | 0.1 to 627.997 | 195 | 35.23 | -394.15 | 5.83E-05 | -2.99 | 0.06 |
| 20MgAl-CFX_T2 | 0.1 to 627.997 | 195 | 98.97 | 23.4 | 10.63 | -0.12 | 0.99 |
| 5ZrP-GTM_T1 | 0.1 to 627.997 | 195 | 779.17 | 48.87 | 1469.42 | 0.35 | 0.99 |
| 5ZrP-GTM_T2 | 0.1 to 627.997 | 195 | 6230.46 | 30.6 | 1992.92 | 0.1 | 0.99 |
| 10ZrP-GTM_T1 | 0.1 to 627.997 | 195 | 1126.36 | 27.6 | 1258.14 | 0.37 | 1 |
| 10ZrP-GTM_T2 | 0.1 to 627.997 | 195 | 106.92 | 30.38 | 17.26 | 0.24 | 0.99 |
| 20ZrP-GTM_T1 | 0.1 to 627.997 | 195 | 258.77 | 23.49 | 684.12 | 0.47 | 0.99 |
| 20ZrP-GTM_T2 | 0.1 to 627.997 | 195 | 2139.64 | 57.65 | 1367.77 | 0.27 | 0.99 |
